# Cytosine methylation dynamics during post-testicular sperm maturation in mammals

**DOI:** 10.1101/2020.02.14.949487

**Authors:** Carolina Galan, Ryan W. Serra, Fengyun Sun, Vera D. Rinaldi, Colin C. Conine, Oliver J. Rando

**Affiliations:** Department of Biochemistry and Molecular Pharmacology, University of Massachusetts Medical School, Worcester, MA 01605, USA; University of Pennsylvania Perelman School of Medicine and Children’s Hospital of Philadelphia, Philadelphia, PA 19104, USA

## Abstract

Beyond the haploid genome, mammalian sperm contribute a payload of epigenetic information which can modulate offspring phenotypes. Recent studies have shown that the small RNA payload of sperm undergoes extensive remodeling during post-testicular maturation in the epididymis. Intriguingly, epididymal maturation has also been linked to changes in the sperm methylome, suggesting that the epididymis might play a broader role in remodeling the sperm epigenome. Here, we build on prior studies of the maturing sperm methylation landscape, further characterizing the genome-wide methylation landscape in seven germ cell populations collected from throughout the male reproductive tract. Overall, we find very few changes in the cytosine methylation landscape between testicular germ cell populations and cauda epididymal sperm, demonstrating that the sperm methylome is largely stable throughout post-testicular maturation. Intriguingly, although our sequencing data suggested that caput epididymal sperm exhibit a highly unusual methylome, follow-up studies revealed that this resulted from contamination of caput sperm by extracellular DNA. Extracellular DNA formed web-like structures that ensnared sperm, was present only in the caput epididymis of virgin males, where it was associated with citrullinated histone H3 and presumably resulted from a PAD-driven genome decondensation process. Taken together, our data emphasize the stability of the cytosine methylation landscape in mammalian sperm, and identify a surprising but transient period during which immature sperm are associated with extracellular DNA.

## INTRODUCTION

In addition to contributing a haploid genome to the next generation, it is increasingly clear that germ cells also deliver epigenetic information to progeny that can impact early development and later phenotypes. In mammals, this is best-characterized in the context of genomic imprinting, a situation where genes exhibit monoallelic expression from either the maternal or paternal allele. Heritable marking of regulatory regions that control imprinted genes relies on the covalently-modified cytosine derivative 5-methylcytosine (Bartolomei and Ferguson-Smith, 2011; Bestor and Bourc’his, 2004; Feng et al., 2010; Hackett and Surani, 2013). In mammals, cytosine methylation typically occurs in the context of CpG dinucleotides, and cytosine methylation patterns are readily copied during S phase by the maintenance methyltransferase DNMT1.

Cytosine methylation patterns in mammals undergo two major reprogramming events, the first during primordial germ cell development, and the second occurring shortly after fertilization. In the zygote, sperm methylation is rapidly erased, apparently via active demethylation, shortly after fertilization, while oocyte methylation patterns are lost more slowly via passive demethylation (replication without maintenance methylation). However, a small subset of genomic loci escape this demethylation process, including imprinting control regions and a subset of evolutionarily-young repeat elements (Law and Jacobsen, 2010; Smith and Meissner, 2013). The mechanisms responsible for protecting these loci from demethylation are still being uncovered, but recent studies implicate two sequence-specific DNA binding proteins (ZFP57 and ZFP445) in maintenance of methylation levels at a subset of imprinting control regions (Shi et al., 2019; Takahashi et al., 2019).

Although the majority of the sperm methylation landscape is erased upon fertilization, the existence of escaper loci suggests the potential for environmental effects on the sperm methylation profile to transmit information to the next generation. Indeed, several studies have documented changes in sperm cytosine methylation in response to various diets or toxin exposures (Holland et al., 2016; Radford et al., 2014; Sun et al., 2018; Wei et al., 2014), raising the question of how the sperm methylome is regulated by environmental conditions.

Intriguingly, an early study on sperm methylation revealed the possibility of a third cycle of sperm methylation reprogramming, occurring during post-testicular sperm maturation in the epididymis (Ariel et al., 1994). Briefly, using methylation-sensitive restriction enzymes, the authors showed that methylation levels at three genomic loci appeared to change as sperm entered the proximal, or caput, epididymis. Given recent findings that the epididymis plays a central role in shaping another epigenetic information carrier, the small RNA payload, in sperm (Nixon et al., 2015; Reilly et al., 2016; Sharma et al., 2016; Sharma et al., 2018), we envisioned the exciting possibility that the epididymis may play a broader role in control of the heritable sperm epigenome.

We therefore revisited the prior study by Ariel *et al*, using the gold standard whole genome bisulfite sequencing (WGBS) to characterize the methylation landscape genome-wide, at single-nucleotide resolution, in seven germ cell populations from primary spermatocytes to vas deferens spermatozoa. We found remarkably consistent methylation profiles in five of the seven germ cell populations, with strong correlations between the various testicular germ cell populations and corpus and cauda epididymal sperm. This general persistence of methylation patterns was interrupted by caput epididymal sperm, which were characterized by modest global hypomethylation accompanied by hypermethylation of germline-associated CpG islands. We ultimately identified cell-free DNA, presumably derived from somatic cells of the epididymis, as the cause of the unusual “caput sperm” methylome – most definitively, we show that treating caput sperm with DNase I prior to sperm lysis restored the caput sperm methylation landscape to the same pattern seen from testicular through cauda epididymal sperm. Cell-free DNA was associated with the citrullinated histone H3 (citH3) that is diagnostic of arginine deimination by PAD enzymes, and, curiously, was detected only in virgin males. Taken together, our data support a static view of stable methylation patterns persisting throughout post-testicular sperm development, and reveal an unexplained but intriguing transient stage of programmed cell-free DNA production in the male reproductive tract.

## RESULTS

### Genome-wide analysis of the sperm cytosine methylation landscape

We set out to build on prior low-throughput studies documenting differences in cytosine methylation between testicular sperm and sperm obtained from various regions of the epididymis (Ariel et al., 1994). To this end, we collected seven germ cell populations from 10-12 week old FVB males: primary spermatocytes, two populations of round spermatids, and spermatozoa obtained from the caput, corpus, and cauda epididymis, and from the vas deferens (**Figure 1A**). For each population we collected samples from seven different males, isolated genomic DNA, and prepared libraries for whole genome bisulfite sequencing (WGBS). Overall we obtained an average of ∼300 million reads per sample, representing a mean coverage of ∼15X per CpG. These data confirmed well-described features of the sperm methylome (Molaro et al., 2011), such as a high overall level of methylation interrupted by hypomethylated CpG islands (**Figures 1B-C**), supporting the quality of our dataset.

**Figure 1.**
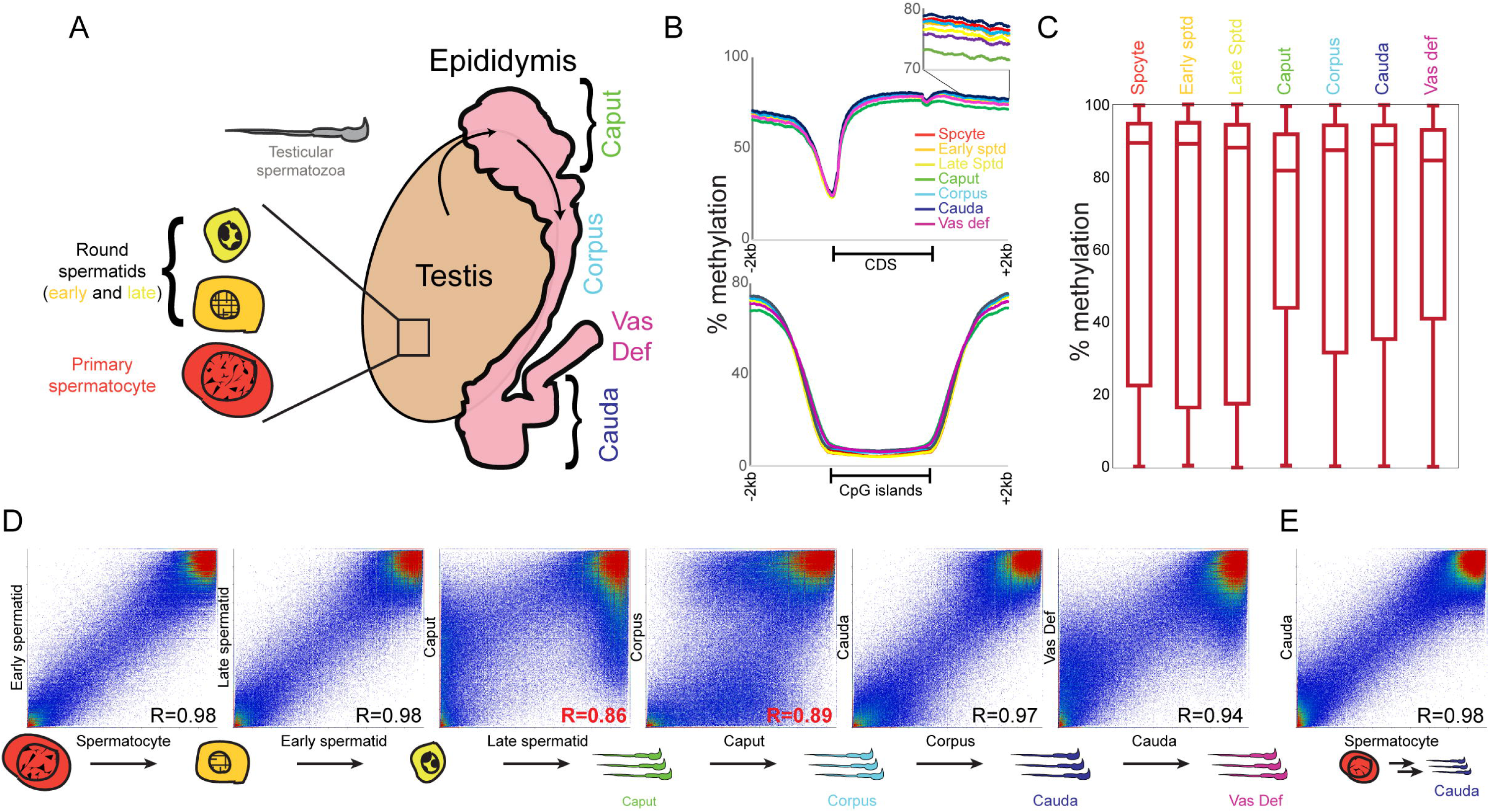
Whole genome cytosine methylation in seven germ cell populations. A) Schematic of seven germ cell populations analyzed in this study: primary spermatocytes, early and late round spermatids from testis, and spermatozoa from the caput, corpus, cauda epididymis and from the vas deferens. Shown in grey are mature testicular spermatozoa, which were not analyzed here. B) Typical features of the germline methylome are reproduced in all seven datasets. Top panel shows average methylation profile for metagenes normalized to the same length, along with 2 kb of sequence upstream and downstream of genes. Bottom panel shows methylation data surrounding all annotated CpG islands. C) Box plots show methylation levels for all 200 bp tiles across the genome. As expected, the majority of samples are overwhelmingly methylated, as previously observed, with caput sperm and vas deferens sperm exhibiting modest hypomethylation relative to the other five samples. D) Scatterplots comparing methylation levels for consecutive sperm developmental stages. In each case, scatterplot shows methylation levels averaged for all 200 bp tiles (with at least 10 methylation-informative reads) across the genome. Overall, all samples exhibit robustly correlated methylation landscapes with one another, with the caput and vas deferens samples representing outliers with hundreds of hypo- and hyper-methylated tiles compared to other sperm samples. See also **Figure1-figure supplement 1** for all pairwise comparisons. E) Scatterplot comparing methylation levels in primary spermatocytes and cauda sperm, showing that the genomic loci that exhibit changes in methylation in caput sperm return to their original methylation levels later in the epididymis.

### A transient methylation signature in the caput epididymis interrupts otherwise stable methylation throughout sperm maturation

We next turned to the question of how the sperm methylome changes over the course of epididymal transit. Comparing all seven populations, we noted similar overall methylation profiles, with global methylation punctuated by hypomethylated CpG islands in all seven samples (**Figures 1B-C**). Intriguingly, we find somewhat lower global methylation in caput sperm and, to a lesser extent, in sperm from the vas deferens. Examination of metagenes revealed that these two sperm populations deviated somewhat from the characteristic methylation profile observed in the other five samples, with lower methylation levels across coding regions, along with moderately increased methylation at promoters (**Figure 1B** and see below).

To more systematically search for methylation changes between germ cell populations, we averaged methylation levels over 200 bp regions tiled across the genome and compared methylation levels in these tiles between all pairs of samples in our dataset (**Figure 1D** and **Figure 1-figure supplement 1**). Overall, we find that five germ cell populations exhibited nearly-identical methylation landscapes, with strong correlations between the methylation datasets for the three testicular samples, and for corpus and cauda sperm. Importantly, this means that the differences in methylation between round spermatids and caput sperm (see below) are reversed in corpus sperm, rather than the round spermatid->caput and caput->corpus methylation changes being part of an ongoing process of progressive methylation maturation in the epididymis. In other words, the nearly-identical methylation profiles for testicular spermatocytes/spermatids and cauda sperm (**Figure 1E**) indicates that the sperm methylome is essentially unchanged by the process of epididymal maturation, and strongly argues against a major role for the epididymis in modulating sperm methylation.

Below, we explore the unusual methylome of caput sperm. For simplicity’s sake, we will generally ignore the similar behavior of vas deferens sperm, but we reconcile caput and vas deferens sperm in later analyses.

### Widespread derangements in CpG island methylation in caput epididymal sperm

In contrast to the nearly-identical methylation profiles obtained from testicular and corpus/cauda sperm, comparisons between caput sperm and any of these samples revealed large-scale changes in methylation across hundreds of 200 bp tiles (**Figure 1D** and **Figure 1-figure supplement 1**). Overall, as noted above, caput sperm were slightly less methylated than these other sperm samples – examination of hypomethylated loci in caput sperm revealed diffuse hypomethylation over a wide range of both coding and intergenic genomic loci (not shown). This signature was also detectable in metagene averages in the highly-methylated regions distant from CpG islands, where caput sperm exhibited a small global deficit at these loci (**Figure 1B**, insets).

In addition to the widespread hypomethylation of the caput sperm genome, a large group of hypermethylated tiles was readily apparent in these scatterplots. Examination of these tiles revealed that they were largely associated with CpG islands. To visualize this, we calculated the average methylation across all annotated CpG islands; **Figure 2A** shows a scatterplot comparing CpG island methylation in round spermatids and caput sperm – nearly-identical results were obtained in comparisons between caput sperm and other testicular populations, or corpus/cauda epididymal sperm. Overall, we find widespread methylation changes at regulatory elements in caput sperm, primarily reflecting hypermethylation of CpG islands in this sample (dots above and to the left of x=y) with a relatively small number of hypomethylated islands in caput sperm. **Figure 2B** extends this analysis to show data for individual islands, showing methylation levels for all seven germline populations relative to the mean methylation value for each CpG island, highlighting the dramatic changes in CpG island methylation in caput sperm (and, to a lesser extent, in vas deferens sperm). Again, we note that regulatory elements that exhibit aberrant methylation in caput sperm universally return to the spermatocyte/spermatid methylation patterns in the corpus and cauda samples. In other words, whatever methylation changes apparently occur in caput sperm are soon reversed in later sections of the epididymis (with the exception of the vas deferens, discussed later).

**Figure 2.**
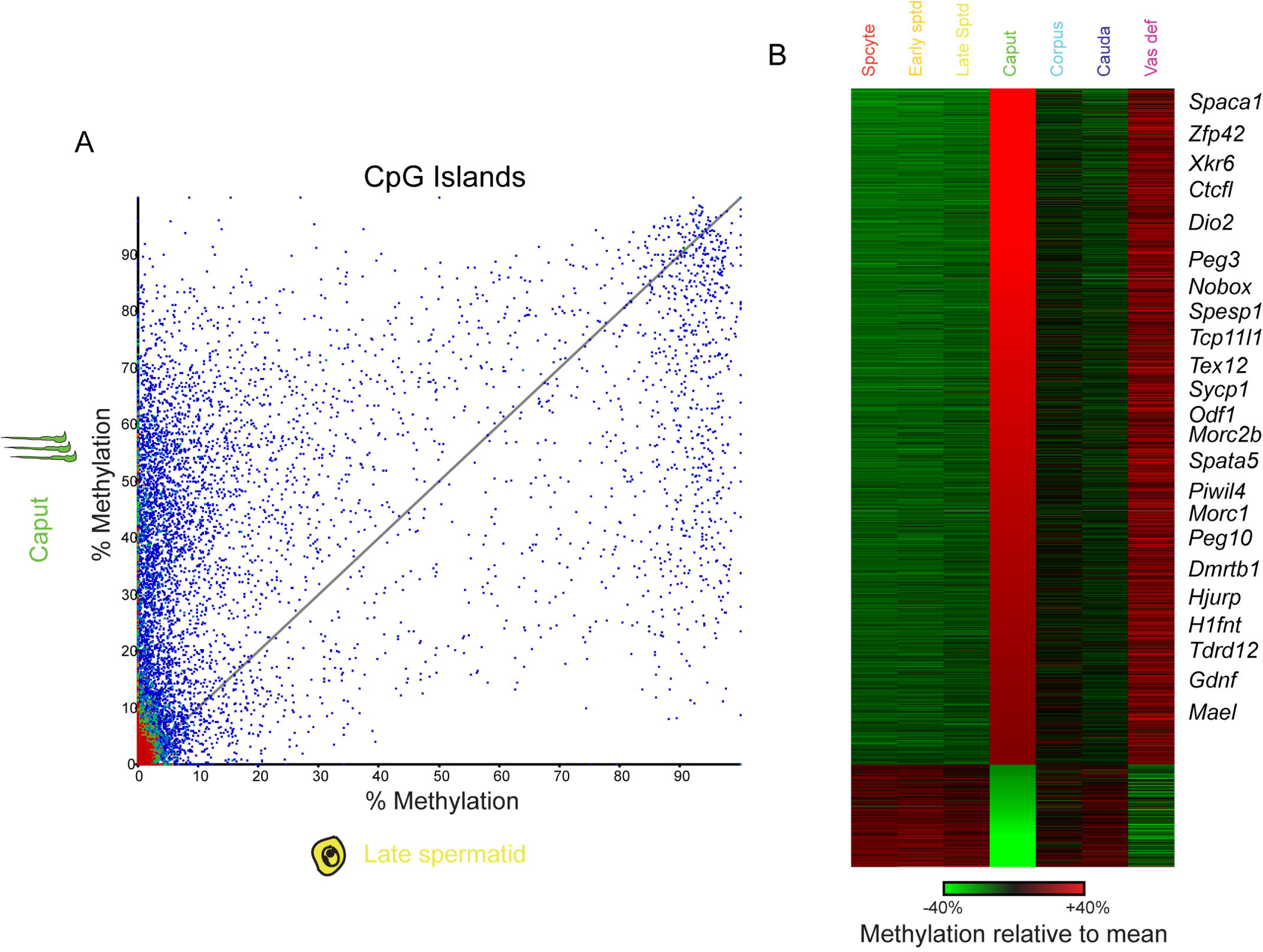
CpG island hypermethylation in caput and vas deferens sperm. A) Scatterplot showing average methylation levels across all annotated CpG islands, comparing late round spermatids and caput epididymal sperm. Although the majority of CpG islands are still hypomethylated in both samples (lower left corner), caput sperm are characterized by hundreds of abnormally hypermethylated CpG islands (above the x=y diagonal). B) Heatmap for CpG islands that are more than 20% hypo- or hyper-methylated in caput sperm relative to other samples. Each row represents the methylation values of a single CpG island, normalized relative to the average methylation % across all seven samples. Islands are sorted from hyper-methylated in caput sperm to hypo-methylated. As caput-hypermethylated CpG islands were enriched for those located near genes with known reproductive functions, notable genes are annotated along the right of the heatmap. We note that the majority of caput-*hypo*methylated CpG islands (bottom of the heatmap) are located within transcribed regions, and may therefore reflect the modest global hypomethylation observed over transcribed regions (**Figures 1B-C**).

What biological pathways are affected by epididymal methylation dynamics? To address this question, we sought gene ontology categories enriched in both caput sperm hypo- and hyper-methylated CpG islands. Hypomethylated islands in caput sperm were significantly enriched for a handful of processes related to neuronal function (neuronal cell body) and transcriptional regulation (positive regulation of DNA-templated transcription). However, these enrichments appear to be primarily driven by the fact that the majority of caput-hypomethylated CpG islands were found within transcribed regions – the hypomethylation at these loci presumably reflects the more general hypomethylation characteristic of caput sperm (**Figure 1B-C**). More intriguingly, *hyper*methylated islands were enriched for a wide variety of annotations associated with meiosis- and sperm-specific functions, including synaptonemal complex, piRNA biogenesis, ion signaling, and reproductive process.

To validate our genome-wide dataset, and to develop a set of cost-effective targets for follow-up mechanistic studies, we analyzed methylation levels at a number of target loci by pyrosequencing of bisulfite-converted DNA. As shown in **Figure 3A** (and in many further examples below), methylation differences between caput and cauda were robustly reproducible in many additional, independent pairs of sperm samples – for several loci, methylation differences were confirmed in ten pairs of caput and cauda sperm samples overall. These data validate our overall dataset, and will be used in the mechanistic follow-up studies described below. Importantly, although we considered the possibility that a signature of hypermethylation of germline regulatory elements might reflect contamination by somatic cells, we ensured that the methylation differences between caput and cauda sperm were robust to several sperm purification protocols including one based on detergent washing of caput epididymis luminal contents, and one based on Percoll-based isolation of caput sperm in the absence of detergent treatment (**Methods**). For all samples we routinely determined, based on the characteristic hook-shaped morphology of the murine sperm head, that our caput sperm preps were >95% free of somatic cell contamination (see below).

**Figure 3.**
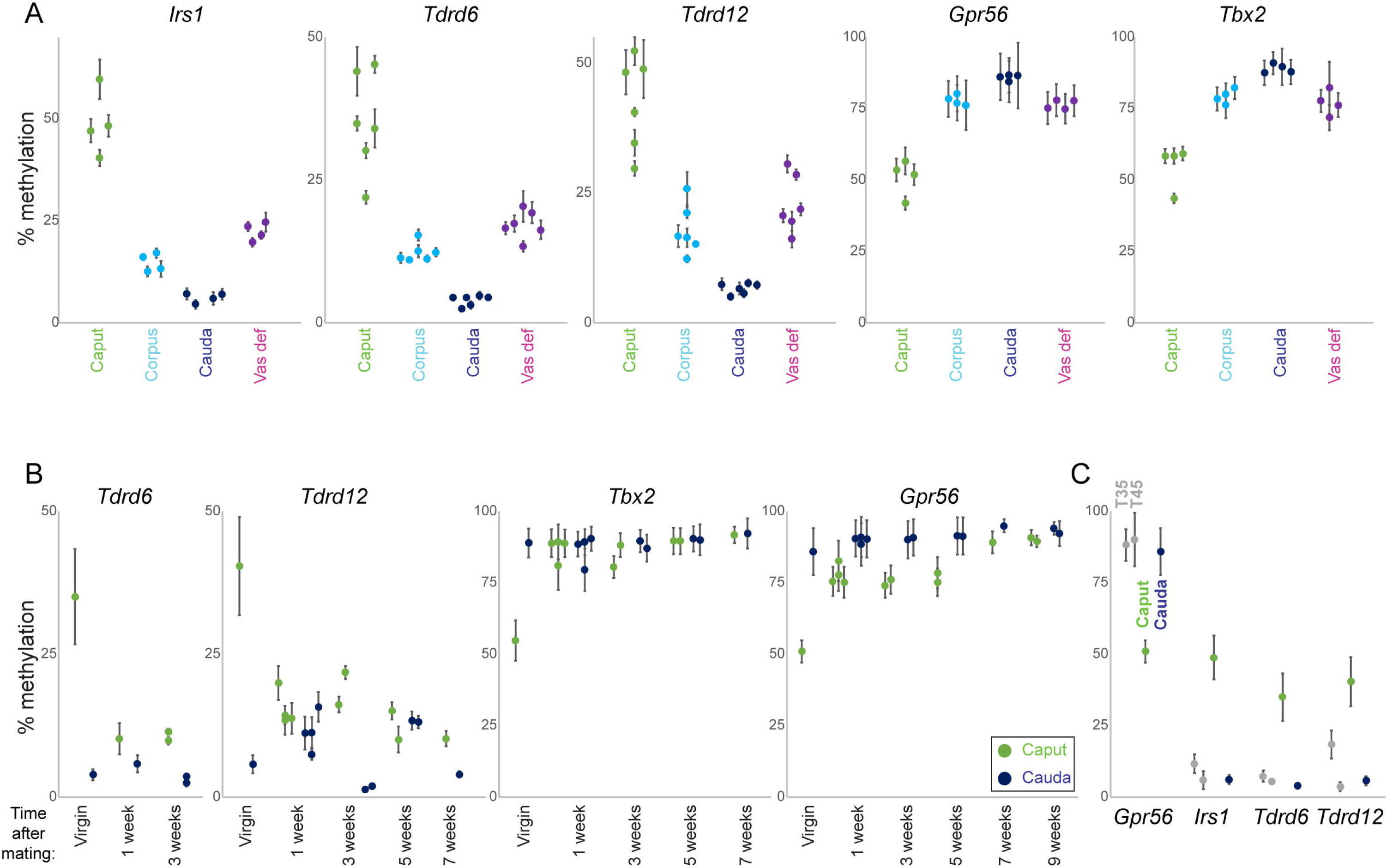
The unusual methylation program in caput sperm is lost after mating. A) Pyrosequencing validation of WGBS data. Pyrosequencing data are shown for five loci which exhibited caput-specific methylation levels in our WGBS dataset. For each locus, pyrosequencing data are shown for 4-5 samples each of caput, corpus, and cauda epididymal sperm, as well as vas deferens sperm, as indicated. Error bars show standard deviation across multiple CpGs assayed at each locus. In all cases the pyrosequencing data recapitulate the differences between sperm populations identified in our WGBS dataset. B) The caput-specific methylation profile is specific to virgin males. Pyrosequencing data shown as in panel (A), for caput and cauda sperm obtained from virgin males or males sacrificed at varying times after confirmed mating. C) Testicular sperm obtained from prepubertal males (thus enriched with the first meiotic wave of spermatogenesis) exhibit similar methylation to cauda, not caput, sperm from mature animals. For each target locus, data are shown for testicular sperm obtained from 35 day or 45 day old males, as well as caput and cauda epididymal sperm obtained from 10-12 week old animals.

### The caput sperm methylome is stable in multiple buffer conditions

We next sought to identify the mechanistic basis for the widespread changes in methylation observed in caput sperm, to explore the molecular basis for what would represent a relatively rare case of fully replication-independent cytosine demethylation. One of the signature functions of the mammalian epididymis is to provide a variety of highly distinctive luminal microenvironments that serve a multitude of functions in supporting sperm maturation and preventing premature activation (Breton et al., 2016; Gervasi and Visconti, 2017). It is well-known that the ionic composition differs extensively between different luminal compartments, as for example pH and calcium levels vary dramatically between caput and cauda, and extensive gene expression differences between different segments (Johnston et al., 2005) suggest that many other metabolites will also differ in concentration throughout the epididymis.

We therefore set out to test the hypothesis that sperm methylation dynamics result from “pre-loaded” genome-associated DNA modification enzymes (DNMTs for methylation, TET enzymes for demethylation) whose ongoing activity could be inhibited or activated by changes in either substrate levels (SAM, alpha-ketoglutarate, iron oxidation status, etc.) or buffer conditions (DNMT3a activity is highly pH-dependent (Holz-Schietinger and Reich, 2015)). To test this, we attempted to recapitulate the caput to cauda methylation changes by incubating purified caput sperm in buffer conditions meant to mimic the cauda epididymal lumen (**Figure 3-figure supplement 1**). However, none of the buffer conditions tested were able to substantially influence the caput sperm methylome.

### The caput sperm methylome is lost following mating

During the course of these studies, we found one animal in which the methylation level at our target genes was nearly-identical for caput and cauda sperm (not shown). Further investigation revealed that a female had accidentally been weaned into an otherwise all-male cage, suggesting that mating might influence the methylation dynamics described here. Indeed, direct testing of this hypothesis confirmed that methylation of our target genes was nearly identical in caput and cauda sperm obtained from animals who had successfully sired offspring (**Figure 3B**). This was due to the caput methylation profile shifting to match that of cauda sperm; cauda sperm methylation was totally unaffected by the male’s mating status.

We considered and rejected two hypotheses regarding the change in methylation status of caput sperm in mated animals: 1) that conversion of the testicular sperm methylome to the unusual caput state might require extended incubation of newly-arrived sperm in the caput luminal environment, and 2) that the virgin caput methylation profile reflects sperm originating from the unusual first meiotic wave of spermatogenesis and somehow being captured in the caput epididymis. The first hypothesis was rejected based on the finding that the cauda-like methylation program in caput sperm from mated animals was stable for at least nine weeks following mating (**Figure 3B**). We next considered the possibility that caput sperm obtained from virgin animals reflect an unusual first wave of spermatogenesis (Grive et al., 2019) characterized by an atypical methylation program. This hypothesis would require an unlikely process – in which first wave sperm would be slowed or arrested in the proximal epididymis, allowing subsequent waves of sperm to pass into the corpus and cauda epididymis – but we could not rule out the hypothesis a priori. However, enriching first wave sperm by isolation of testicular spermatozoa from 35 day-old animals revealed the same methylation levels at our validation loci as those observed for testicular and cauda sperm from older animals (**Figure 3C**), refuting the hypothesis that first wave sperm carry an unusual methylome.

### Contamination of caput epididymal sperm by cell-free DNA

What then could account for the unique methylation program observed in caput sperm from virginal males? Careful inspection of our genomic DNA preparations revealed that although testicular and cauda sperm samples were characterized by uniformly high molecular weight genomic DNA, there was a dim but notable additional “cloud” of lower molecular weight DNA in the caput sperm preps (not shown). This suggested the possibility that caput sperm might be contaminated by cell-free DNA. Indeed, DAPI staining of caput sperm preparations revealed not only the expected hook-shaped sperm heads, but at higher exposure times we noted the presence of web-like DNA structures, often ensnaring multiple sperm (**Figure 4A**). These webs were not detected in cauda epididymal sperm preparations (**Figure 4-figure supplement 1**). To visualize this extracellular DNA in situ, we stained histological sections of several epididymal regions with DAPI. Intriguingly, in both caput sections and in the vas deferens we find a DAPI-staining rim associated with the apical region of the epithelium (**Figure 4B**, rim indicated in yellow). DAPI-positive rims were not observed in the cauda epididymis, and were lost from the caput epididymis and vas deferens following mating (**Figure 4B**, bottom panels). To definitively test whether this extracellular DNA is responsible for the methylation profile of virginal caput sperm, we treated virgin caput sperm with DNase I to eliminate any extracellular DNA prior to extraction of genomic DNA for pyrosequencing analysis. Remarkably, DNase I treatment completely restores the caput sperm methylome to the methylation levels observed in testicular or cauda epididymal sperm (**Figure 4C**), demonstrating that contaminating cell-free DNA present in the virgin caput epididymis is responsible for the unusual caput methylome observed in virgins.

**Figure 4.**
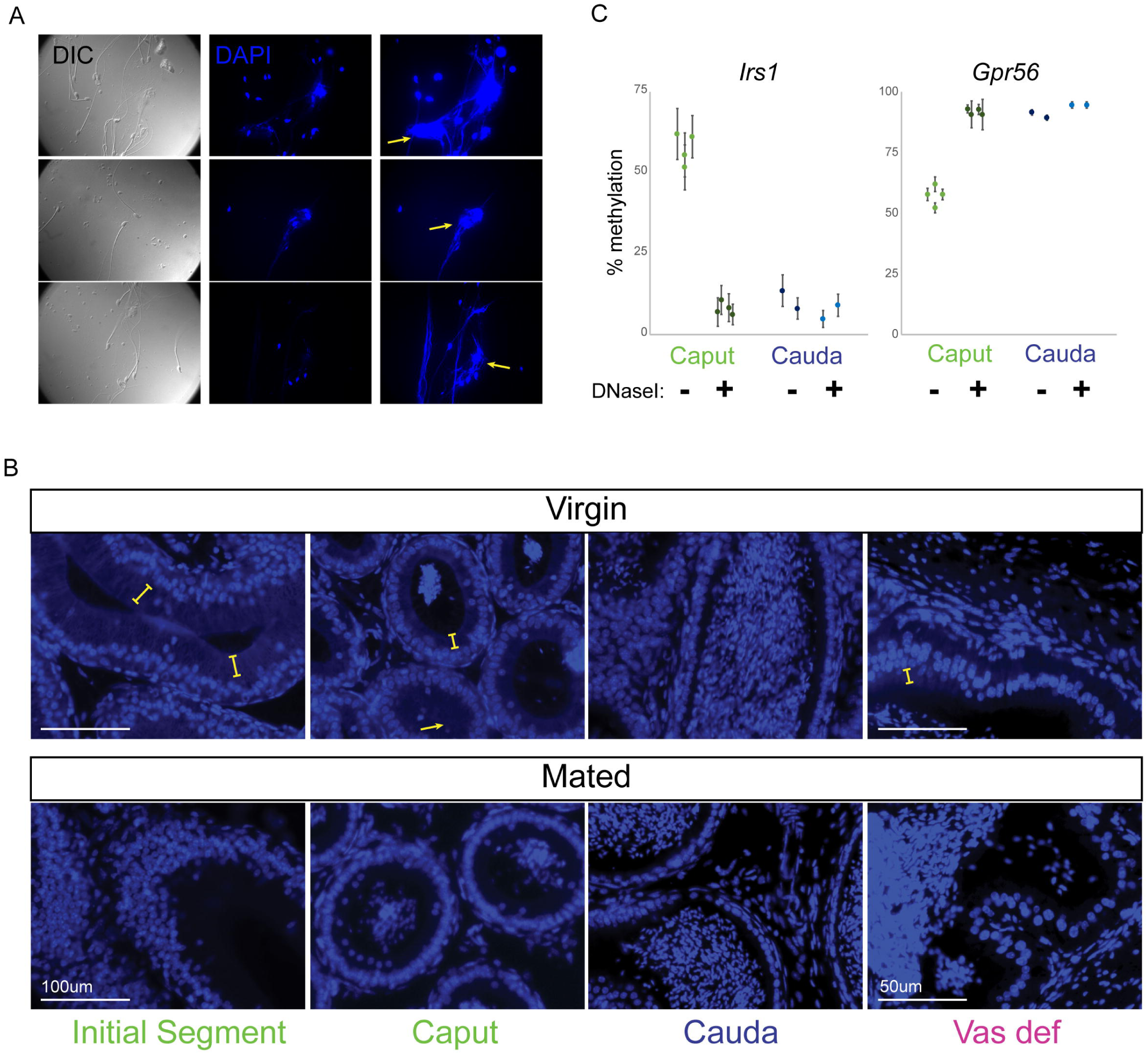
Cell-free DNA in the caput epididymis of virgin males is responsible for the caput methylome. A) DAPI staining of caput sperm samples (See **Figure4-figure supplement 1** for cauda sperm samples). For each sample, left panel shows DIC to show sperm locations, middle and right panels show DAPI staining, with far right panel showing longer exposures. Yellow arrows indicate examples of web-like cell-free DNA in caput sperm samples. B) Cell-free DNA is present in situ in the epididymis. Images show DAPI-stained histology sections taken from the initial segment, caput epididymis, cauda epididymis, and vas deferens of either virgin or mated males, as indicated. Yellow brackets indicate examples of DAPI-positive rim associated with the apical regions of the epithelium of initial segment or caput samples taken from virgin males. Arrow shows a lumen nearly completely filled with this material. C) Cell-free DNA is responsible for the abnormal caput methylome. Pyrosequencing data are shown for caput and cauda sperm samples, either mock treated or pre-treated with DNase I prior to genomic DNA purification.

Based on its impact on caput sperm methylation, it seems likely that the cell-free DNA originates from somatic cells rather than germ cells (see **Discussion**). We noted that the cell-free DNA observed in our sperm preparations are reminiscent of extracellular DNA released by neutrophils, known as Neutrophil Extracellular Traps, or NETs (Brinkmann et al., 2004; Sollberger et al., 2018; Tilvawala and Thompson, 2019). The process of NETosis is associated with increased peptidyl arginine deiminase (PAD) activity, resulting in arginine deimination on histones and other proteins, leaving behind citrulline and leading to massive chromatin decondensation. Examination of RNA-Seq data from throughout the epididymis (Rinaldi et al., 2020) confirmed expression of *Padi2* in this tissue, largely confined to principal cells of the caput epididymis and vas deferens (**Figure 5A**). Moreover, consistent with the caput cell-free DNA being produced downstream of PAD-dependent deimination, we find robust citH3 staining of our caput sperm preparations, with negligible signal in cauda sperm preps (**Figure 5B**).

**Figure 5.**
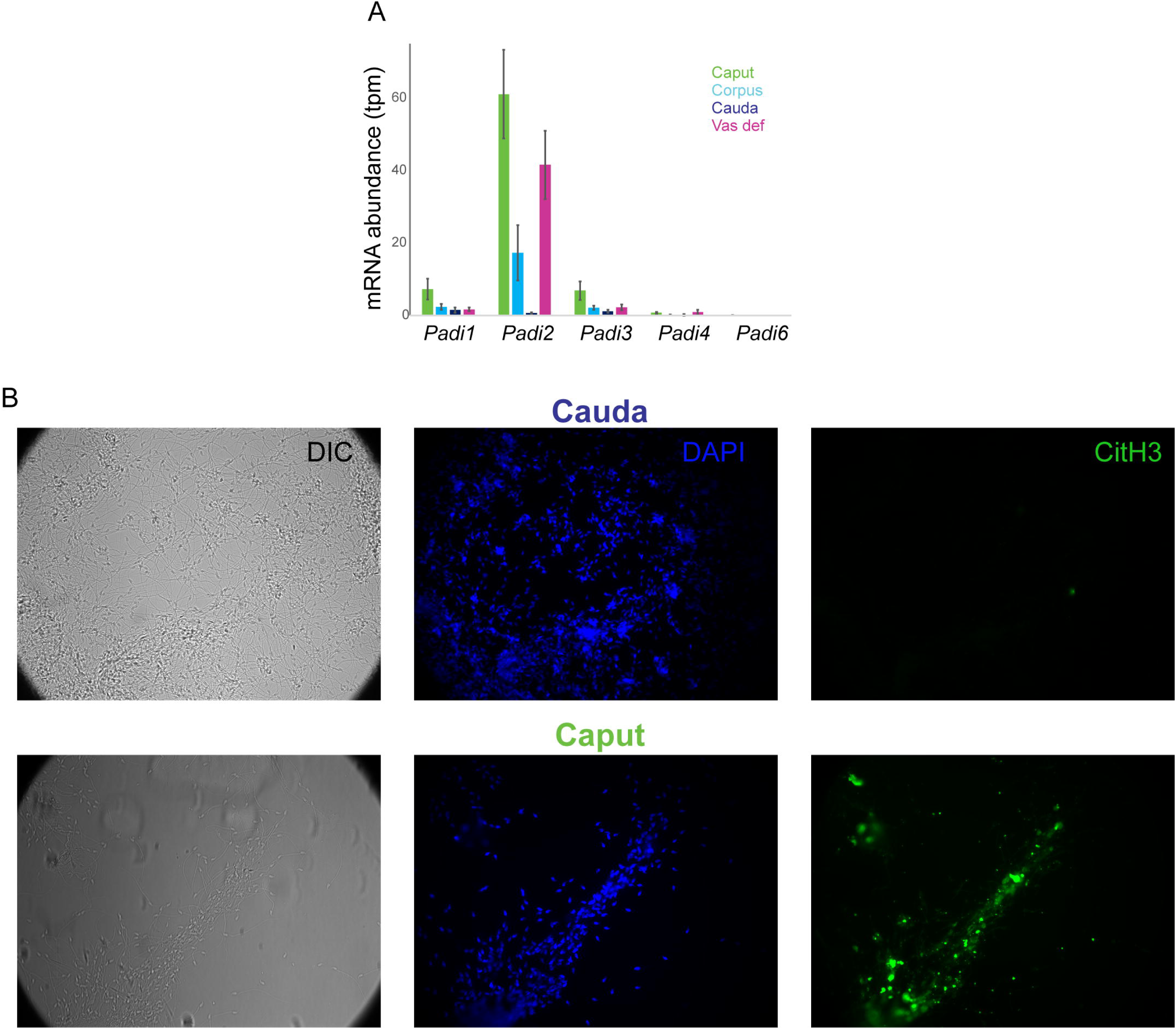
Cell-free DNA is produced via a NETosis-like process. A) Expression of PAD-encoding genes in the murine epididymis and vas deferens. RNA-Seq data are from (Rinaldi et al., 2020). B) Caput epididymal cell-free DNA is associated with citrullinated histone H3. Caput and cauda sperm samples were stained for DAPI (blue) and for citH3 (green), as indicated.

Taken together, our data reveal that PAD-dependent production of extracellular DNA occurs in the epididymis of young males, and that this cell-free DNA is lost following mating.

## DISCUSSION

Here, we characterized the cytosine methylation landscape throughout late stages of spermatogenesis, and throughout post-testicular sperm maturation in the epididymis. Most importantly, these data reveal that the sperm methylome is largely static throughout the process of post-testicular maturation. This is clearest in **Figure 1E**, which shows the extremely robust correlation between the methylation patterns in primary spermatocytes and in mature cauda epididymal sperm. Our data therefore do not support a model in which sperm methylation is extensively remodeled during epididymal transit. Although we cannot rule out subtle methylation changes occurring in epididymal sperm in response to environmental stressors, the absence of changes in this dataset strongly support the classic view of germline cytosine methylation patterns as largely static in the absence of replication-coupled remodeling events.

Our study also revealed a surprising feature of the male reproductive tract, in which cell-free DNA is produced in young virgin males in the caput epididymis and the vas deferens. This DNA almost certainly derives from somatic cells rather than dying sperm, since the methylation profile of virgin caput sperm departs from the characteristic germline methylome to incorporate aspects of the methylation program typical of somatic cells. This is apparent in the hypermethylation of key germline regulatory elements, which are unmethylated and active during spermatogenesis but methylated and repressed in most somatic tissues. This is also clear at imprinting control regions, which are either completely methylated or unmethylated in the sperm genome, but are 50% methylated in somatic cells – the contaminated caput methylome exhibits a shift towards 50% methylation at many imprinting control regions. It is otherwise unclear which cells are responsible for production of cell-free DNA, although it seems most likely that this DNA originates in principal cells of the epididymal epithelium. This is based on 1) our inability to find evidence for neutrophils in a single-cell atlas of the epididymis (Rinaldi et al., 2020); 2) the expression of *Padi2*, rather than *Padi4* which is more typically associated with NETosis, in our RNA-Seq dataset (**Figure 5A**); 3) inability to detect substantial staining of the neutrophil marker myeloperoxidase in our cell-free DNA (not shown); and 4) expression of *Padi2* specifically in principal cells in our single-cell atlas of the epididymis (Rinaldi et al., 2020).

Whatever the cell of origin of this extracellular DNA, it seems likely that it is produced via a PAD-dependent genome decondensation event analogous to the one that drives the production of Neutrophil Extracellular DNA Traps, or NETs (Sollberger et al., 2018). This hypothesis is motivated by the expression of *Padi2* specifically in the two regions exhibiting cell-free DNA, and by the confirmed presence of abundant citH3 staining in caput sperm preps (**Figure 5**). What is the function of the extracellular DNA in the epididymal lumen? Extracellular DNA plays a key role in defense against pathogens in several contexts – NETs in the mammalian bloodstream entangle pathogens and aid in their engulfment by antigen-processing cells (Sollberger et al., 2018), while extracellular DNA in pea (*P. sativum*) roots was shown to protect against fungal infections (Wen et al., 2009). While a similar role could be imagined for epididymal DNA helping to prevent ascending infections, the fact that this material is present only in virgins but lost after mating does not support this idea, as infections would presumably be more likely in sexually-active animals than in virgins. The simplest hypothesis then would be that the extracellular DNA documented here is a vestige of some earlier developmental process – death of epithelial cells during epididymal morphogenesis, for instance – and serves no biological function. That said, the use of PAD-dependent cell death would be unusual for typical programmed cell death in development. Alternatively, we speculate that the DNA webs observed here could ensnare sperm rather than pathogens, potentially playing a role in removal of early rounds of defective sperm or some similar process.

Although the function, if any, served by luminal extracellular DNA is unclear, our findings do raise two important technical considerations for reproductive biology studies. First, one important implication of this finding is that at least some aspects of sperm populations are different between virgins and mated males, raising the question of whether other features of the epididymal sperm epigenome – often measured in virgin males – are stable throughout reproductive life. For example, we and others have previously documented substantial differences between the small RNAs carried by caput and cauda epididymal sperm, with a variety of genomically-clustered microRNAs, and a subset of tRNA fragments such as tRF-Val-CAC, being far more abundant in cauda sperm than in caput (Nixon et al., 2015; Sharma et al., 2016; Sharma et al., 2018). We therefore repeated small RNA-Seq for caput and cauda sperm samples obtained from males three or six weeks after confirmed mating. Importantly, we confirm the same overall differences between these sperm samples in their small RNA payload (**Figure 5-figure supplement 1**), demonstrating that at least this observation is not an artifact of some transient early process in virgins. That said, other aspects of the sperm maturation process may differ between virgins and mated males, emphasizing the importance for reproduction studies to explicitly state the mating status of males being used.

The other technical implication of our work regards studies of the sperm methylome, as future efforts focused on cytosine methylation in sperm must contend with the possibility of contamination by cell-free DNA. Although most such studies do not utilize caput epididymal sperm, it is well-known that different labs differ in whether “mature sperm” used for molecular studies or IVF are isolated only from the cauda epididymis, or a mixture of cauda and vas deferens. Evidently, dissections that capture variable lengths of the vas deferens will result in variability in sperm methylation due to the contamination of these samples by DNA with a somatic cell methylation program. In addition, although we find contaminating DNA specifically in the caput and vas of virgins, it is plausible that cell-free DNA could be more persistent or produced in different parts of the epididymis in other strain backgrounds or in males subject to different diets or stressors. A simple solution to this issue of course would be to treat all sperm samples with DNase I prior to any methylation analyses.

## ACKNOWLEDGEMENTS

We thank F. Krueger and S. Andrews (Babraham Institute) for assistance with WGBS analysis, the P. Thompson lab for generous gift of antibodies (citH3 ab5103) and for reading the manuscript, the N. Rhind lab for assistance with microscopy, and A. Boskovic and other members of the Rando lab for insightful discussions. This work was supported by NIH grants R01HD080224 (OJR) and F31HD097928 (CG).

## METHODS

### Mice

Tissues were obtained from 10-12 week old male FVB/NJ mice. All animal care and use procedures were in accordance with guidelines of the University of Massachusetts Medical School Institutional Animal Care and Use Committee.

### Dissection and sperm purification

FVB mice, euthanized at 10-12 weeks of age (unless otherwise noted) according to IACUC protocol, were dissected into four segments that roughly corresponded to caput, corpus, cauda, and vas deferens. Caput, corpus, and cauda epididymis as well as vas deferens were placed into Donners complete media (Hisano et al., 2013) and tissue was cleared of fat and connective tissue before incisions were made using a 26G needle while keeping the bulk tissue intact. Tissue was gently squeezed allowing sperm to escape into solution. After incubation at 37°C for 1 hour, sperm containing media was transferred to a fresh tube and collected by centrifugation at 5000rpm for 5 minutes followed by a 1X PBS wash. To eliminate somatic cell contamination, sperm were subjected to a 1mL 1% Triton X-100 incubation 37°C for 15 mins with 1500 rpm on Thermomixer and collected by centrifugation at 5000rpm for 5 minutes. Somatic cell lysis was followed by a 1x ddH2O wash and 30 second spin 14000 rpm to pellet sperm.

### Isolation of caput sperm using a discontinuous Percoll gradient

Sperm collection from the caput epididymis is performed as described above, but without somatic cell lysis. Instead, caput sperm are purified using a Percoll gradient as described (Krapf et al., 2012) to separate somatic cells away from sperm. Briefly, the caput sperm suspension is carefully layered over a discontinous Percoll gradient containing 45% Percoll (upper phase) and 90% Percoll (lower phase). After centrifugation for 25 min at room temperature (650xg), the interphase containing caput sperm is washed with 1x PBS and prepped for downstream analysis.

### Testicular spermatocyte and spermatid isolation

For each isolation, two testes were acquired from one FVB/NJ mouse at 10-12 weeks of age. Cell suspension was prepared by incubating the testes without their tunica albuginea in 5 ml elutriation buffer (100 mM NaCl, 45 mM KCl, 6 mM Na_2_HPO_4_, 0.6 mM KH_2_PO_4_, 0.23% Sodium DL-Lactate, 0.1% Glucose, 0.1% BSA, 0.011% Sodium Pyruvate, 1.2 mM MgSO_4_ and 1.2 mM CaCl_2_) containing 25 µg/ml liberase TM (Roche Diagnostics GmbH) for 30 min at 37 °C with gentle agitation every 5 min. The cell suspension was mixed by pipetting 20 times with a 10-ml plastic pipette. After homogenization by pipetting 10 times through a P1000 pipette, the single cell suspension was filtered twice through a 40-μm cell strainer (Fisher Scientific) on ice and centrifuged at 1500 rpm at 4 °C for 10 min, and then the pellet was resuspended with 20 ml elutriation buffer. Separation of testis cell populations was performed by centrifugal elutriation using a JE-5.0 elutriation system and a 4-ml standard elutriation chamber (Beckman Coulter). The assembly of the system followed the manufacturer’s instruction. The precise elutriation conditions are as follows: Fractions 1-3 were run at 3000 rpm and fractions 4-5 were run at 2000 rpm. The flow rate was 14, 18, 31, 23, and 40 ml/min for fractions 1-5, respectively. During elutriation, the elutriation chamber was maintained at 4 °C and the cells were collected into 50-ml conical polypropylene tubes that were packed on ice. Cells in tubes of fractions 3 to 5 were pelleted by centrifugation at 1500 rpm at 4 °C for 10 min. All pellets from the same fraction were combined and resuspended in 200 µl of elutriation buffer. Percoll (Sigma-Aldrich) gradient (23-35%) was prepared by using a Gradient Master 108 (Biocomp) following standard program: S1/1, 2:26 (time), 82.0 (angle), 13 (rpm). After loading the cell suspension, centrifugation was carried out with SW40Ti Rotor (Beckman Coulter) at 11,000 rpm at 4 °C for 15 min. The cells were then collected in 15-ml conical polypropylene tubes and pelleted by centrifugation at 1500 rpm at 4 °C for 15 min. The cell pellets were stored at -80 °C and ready for DNA extraction. Based on cell morphology and small RNA data, Fraction 5 was determined to correspond to primary spermatocytes, with fractions 4 and 3 being relatively early and late round spermatids, respectively.

For isolation of the first wave of testicular spermatozoa from postnatal day 35, a cell suspension was prepared by dissecting testes from postnatal day 35 males into 35mm dish containing 150 mM NaCl, removing tunica albuginea, and dissociating tissue using 22G needle and 3 mL syringe with 150 mM NaCl. Cell suspension was then pipetted into a 15 mL conical tube to allow for large tissue to settle. Once settled, top 1mL was loaded onto 52% isotonic Percoll solution. Samples were ultracentrifuged at 15,000 rpm for 10 min at 10 °C. Following ultracentrifugation, pellet was carefully isolated, washed with 150 mM NaCl to remove residual Percoll, resuspended in PBS, and assessed for purity using microscopy (see Figure S1A in (Sharma et al., 2018) for typical preparation) before undergoing the genomic DNA isolation protocol.

### Genomic DNA isolation

Briefly, 750 μL Extraction Buffer (4.24M Guanidine Thiocyanate, 100 mM NaCl, 1% N-Laurylsarcosine, 150mM freshly prepared DTT, 200 μg/mL Proteinase K) was added to sperm pellets and incubated for 2 hours at 56°C with shaking. Samples were then allowed to equilibrate to room temperature after which 600 μL isopropanol was added to precipitate DNA followed by a 15 mins spin at 14,000rpm. Supernatant was carefully discarded and pellets were washed 2x with 80% EtOH, dried briefly, and resuspended in 10 mM Tris pH8 (500 μL) with 5 μL RNase A (Qiagen 19101) for 2 hours after which samples were treated with 5 μL Proteinase K (Qiagen 19131) overnight. Following phenol chloroform extraction (UltraPure Phenol:Chloroform:Isoamyl Alcohol ThermoFisher 15593031) using phase lock gel tubes (Quantabio 10847-802), aqueous phase was transferred to a fresh tube containing 1.5 μL glycogen (20 mg/mL), 1.3 μL 5 M NaCl, gDNA, and 100% EtOH. Genomic DNA integrity was determined using NanoDrop and quantified for downstream applications using Qubit (DNA BR cat Q32850)

### Whole genome bisulfite sequencing

For each population we collected sperm samples from seven different males, isolated genomic DNA, and prepared libraries for whole genome bisulfite sequencing (WGBS); Bisulfite converted DNA was prepared according using the EZ DNA Methylation-Lightning Kit (Zymo D5030). Library construction was performed using TruSeq DNA Methylation Kit (Illumina) and Accel-NGS Methyl-Seq DNA Library Kit (Swift) according to the manufacturer’s instructions. For TruSeq (Illumina) libraries, 50ng bisulfite converted DNA was used and individual libraries were barcoded using the TruSeq DNA Methylation Index PCR Primers (IIlumina) to allow for multiplexed sequencing of libraries. Size selection and DNA purification was performed using Ampure XP beads (Agencourt) according to the TruSeq DNA Methylation kit protocol. For Accel-NGS Methyl-Seq (Swift) libraries, 30ng bisulfite converted DNA was used and barcoded following the (Swift) manufacturer’s instructions including 6 rounds of PCR. All final libraries were analyzed by Fragment Analyzer and quantitated using the Qubit as well as the KAPA Library Quantitation qPCR kit.

### Data analysis

Briefly, sequences were trimmed with Trim Galore (v0.4.4; Cutadapt v1.9.1) - Swift libraries were trimmed by 10bp from their 5’ ends for both R1 and R2 and Truseq libraries were trimmed by 7 bp from their 5’ ends respectively - to avoid biases and mapping issues. The non-CG methylation levels were consistently very low (Swift: 0.2%, Truseq: 0.3-0.6%) indicating good bisulfite conversion rates (>= 99.4%). The resulting trimmed sequences were mapped to the mouse GRCm38 genome using Bismark (Krueger and Andrews, 2011) (v0.17.0); CpG methylation calls were extracted and analysed using SeqMonk (www.bioinformatics.babraham.ac.uk/projects/seqmonk/). All analyses presented in the manuscript were carried out using the Swift dataset (which exhibits better coverage of CpG islands, among other advantages), but qualitatively similar findings were obtained with the Truseq dataset.

### Data availability

Data will be publicly available at GEO, Accession #GSE100220.

### Pyrosequencing

Pyrosequencing was performed using the PyroMark Q24 (Qiagen: 9001514) according to the manufacturer’s instructions. Briefly, bisulfite converted DNA was prepared using the EZ DNA Methylation-Lightning Kit (Zymo D5030). PCR was performed with 10-20ng of bisulfite converted material for each locus of interest (see **Table S1**). Primers (IDT) were designed using the PyroMark Assay Design software (Qiagen)

### Ex vivo sperm incubations

Prior to genomic DNA extraction, sperm were incubated for 4 hours at 37°C in the following conditions: Donners complete media (Hisano et al., 2013), Donners complete media supplemented with either 200 μM SAM, additional sodium pyruvate (10 mM), additional sodium _DL_-Lactate (1% vol/vol), NaHCO_3_ (200mM) as well as Donners complete buffered to pH 6.5, 7, and 7.4. Additional conditions include: Donners basic (stock solution) (Hisano et al., 2013), Donners basic supplemented with NaHCO_3_ (final 25mM), Donners basic supplemented with BSA (final 20 mg/mL), Donners basic supplemented with Sodium _DL_-Lactate (final 0.53% vol/vol), and finally Donners basic buffered to pH 6.5, 7, and 7.4.

DNase treatment of caput sperm was performed prior to gDNA isolation as per manufacturer’s instructions (Qiagen79254)

### Visualization of cell-free DNA and immunostaining

After sperm prep, samples were dried onto VWR® Superfrost® Plus Micro Slides (48311-703), fixed for 10 minutes in 4% paraformaldehyde solution, washed 3x with PBS, and permeabilized for 10 min with 0.1% triton in PBS. Blocking was performed with 10% BSA in PBS for 1 hour at room temperature in a humidified chamber. Staining with primary antibody, Anti-Histone H3 (citrulline R2 + R8 + R17 (Abcam ab5103), was performed for 2 hours at room temperature in 1% BSA PBSt at a concentration of 1:250 in a humidified chamber. Primary antibody was decanted and slides washed three times with PBS for 5 minutes each. Goat anti-Rabbit 488 secondary antibody (Invitrogen A11008) was performed for 1 hour at room temperature at a concentration of 1:500 in a humidified chamber. Secondary antibody was decanted and slides washed three times with PBS for 5 minutes each followed by VECTASHIELD® Antifade Mounting Medium with DAPI (Vector Labs H-1200), sealing coverslip, and imaging on the Zeiss Axioskop 2 Plus.

### Histology

Virgin or retired breeders of approximately same age were anesthetized and perfused with phosphate-buffered saline (PBS) followed by 4% paraformaldehyde (PFA)/PBS. Epididymides were explanted and further incubated in 4%PFA/PBS at 4 °C overnight. After washing the excess of PFA with PBS, the sample was incubated at 4 °C with a 30% sucrose, 0.002% sodium azide in PBS until organ sank to the bottom of the tube. The sucrose solution was replaced and once organ remained at the bottom of the tube the same volume of optimal cutting temperature compound (OCT) was added to the vial and kept ON at 4C under agitation. Samples were than mounted in OCT and frozen at -80 °C until sectioning. Sectioning was done at a thickness of 5 μm by the UMASS morphology core. Slides were store frozen at -20 °C.

Slides were placed at a 37 °C warm plate for 10 minutes to ensure proper attachment of the section to the slide, then washed three times for 5 minutes in PBS 0.02% tween 20 (PBS-T) to remove OCT, followed by DAPI stain (3 μM 4′,6-diamidino-2-phenylindole in PBS) for 5 min. After washing off the excess of DAPI with PBS-T, slides were mounted with ProLong gold Antifade (Thermofisher P36930) and imaged the following day.

### Small RNA Sequencing

Males from the same litter were split into the following groups: not mated, mated and recovered for three weeks, and, mated and recovered for six weeks. Mating was confirmed by the formation of blastocysts in culture. All males were dissected at the same age (14 weeks). Caput and cauda sperm were isolated as described and small RNA sequencing was performed. Isolation of 18–40 nts small RNAs was carried out as previously described (Sharma et al., 2018). Briefly, size selection by purification of RNAs from 15% polyacrylamide-7M urea denaturing gels and sequencing library preparation using Illumina’s TruSeq Small RNA Library Preparation Kit.

## SUPPLEMENTARY INFORMATION

### SUPPLEMENTARY TABLES

**Table S1.**
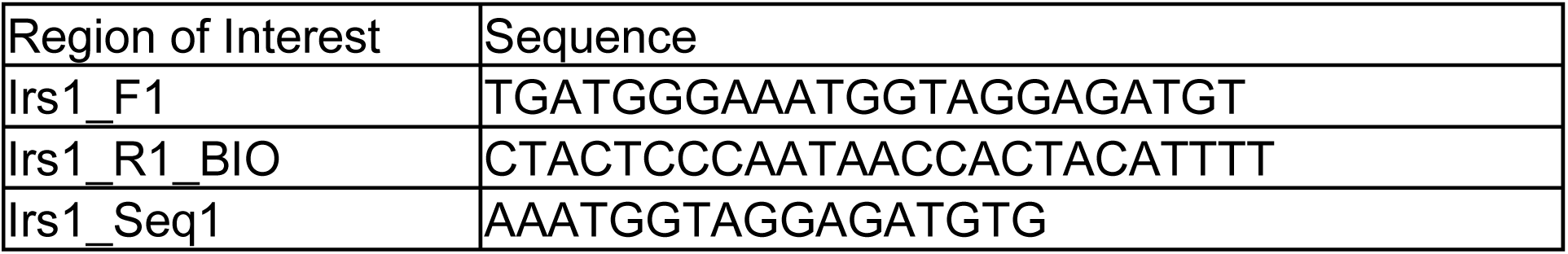
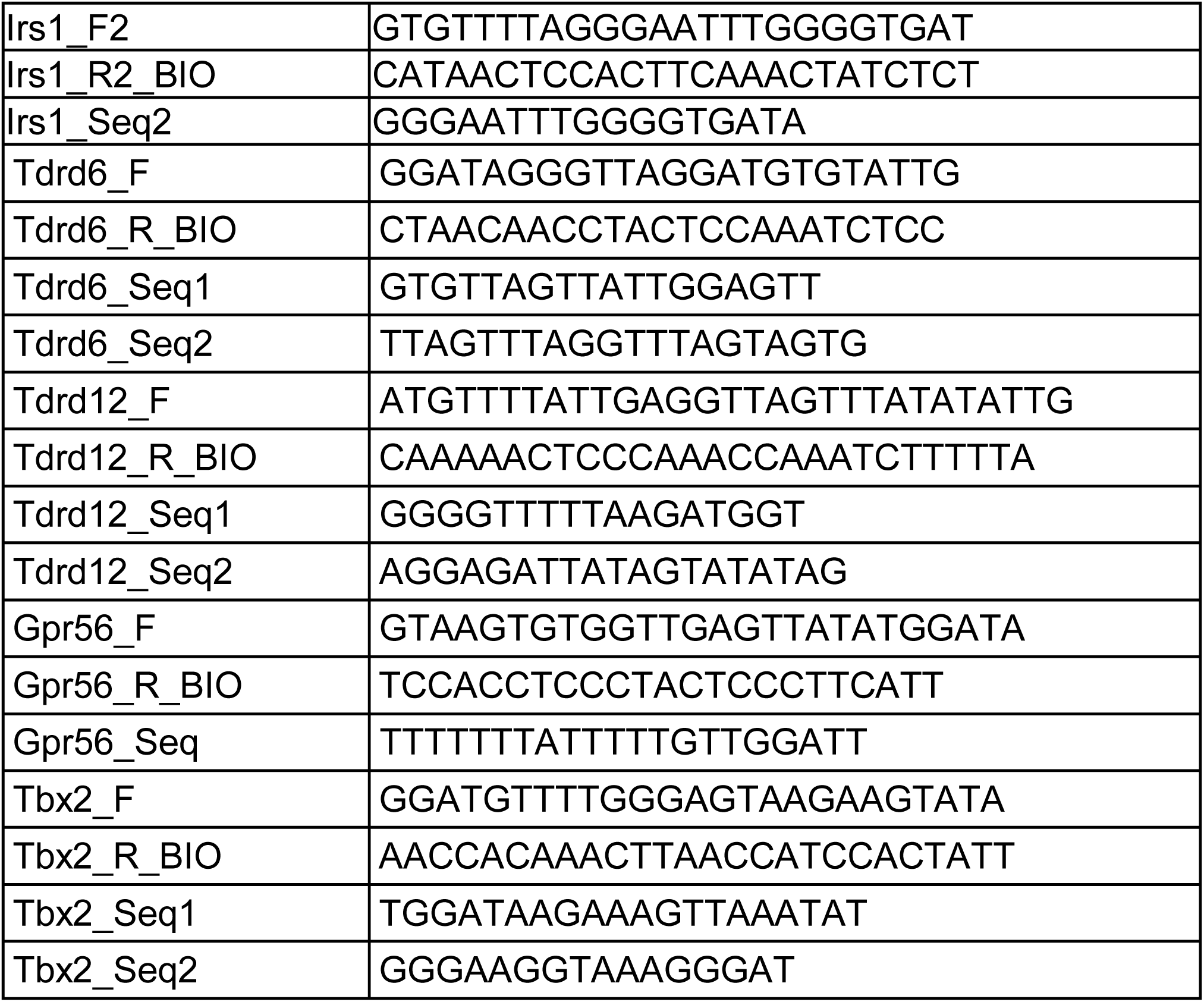
Oligonucleotides used for pyrosequencing. Sequences for pyrosequencing primers used in **Figures 3-4**.

### SUPPLEMENTARY FIGURES

**Figure 1-figure supplement 1.**
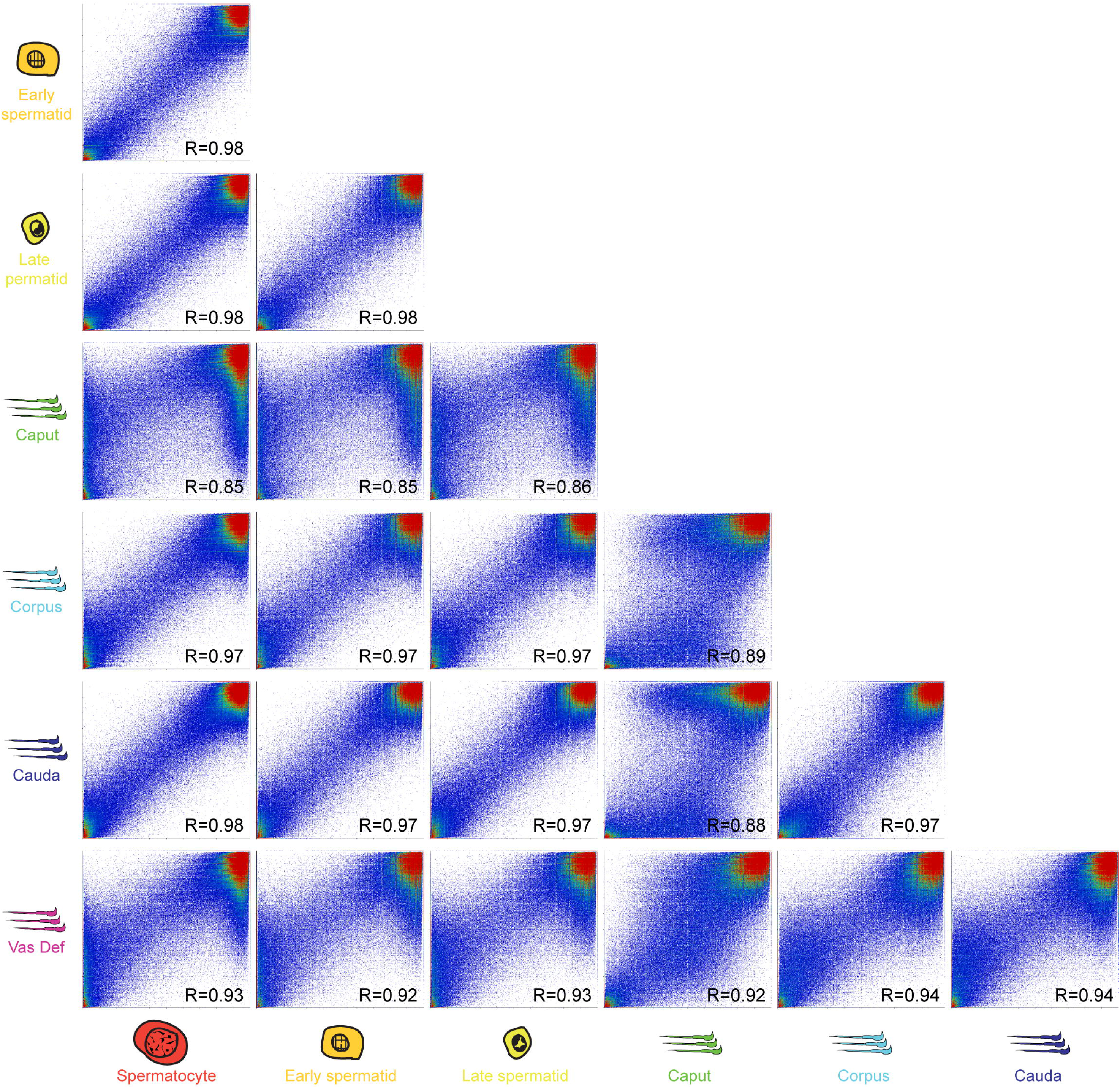
Correlations between all seven germ cell methylation datasets. Scatterplots are shown as in **Figures 1D-E**, for all pairwise comparisons in this dataset.

**Figure 3-figure supplement 1.**
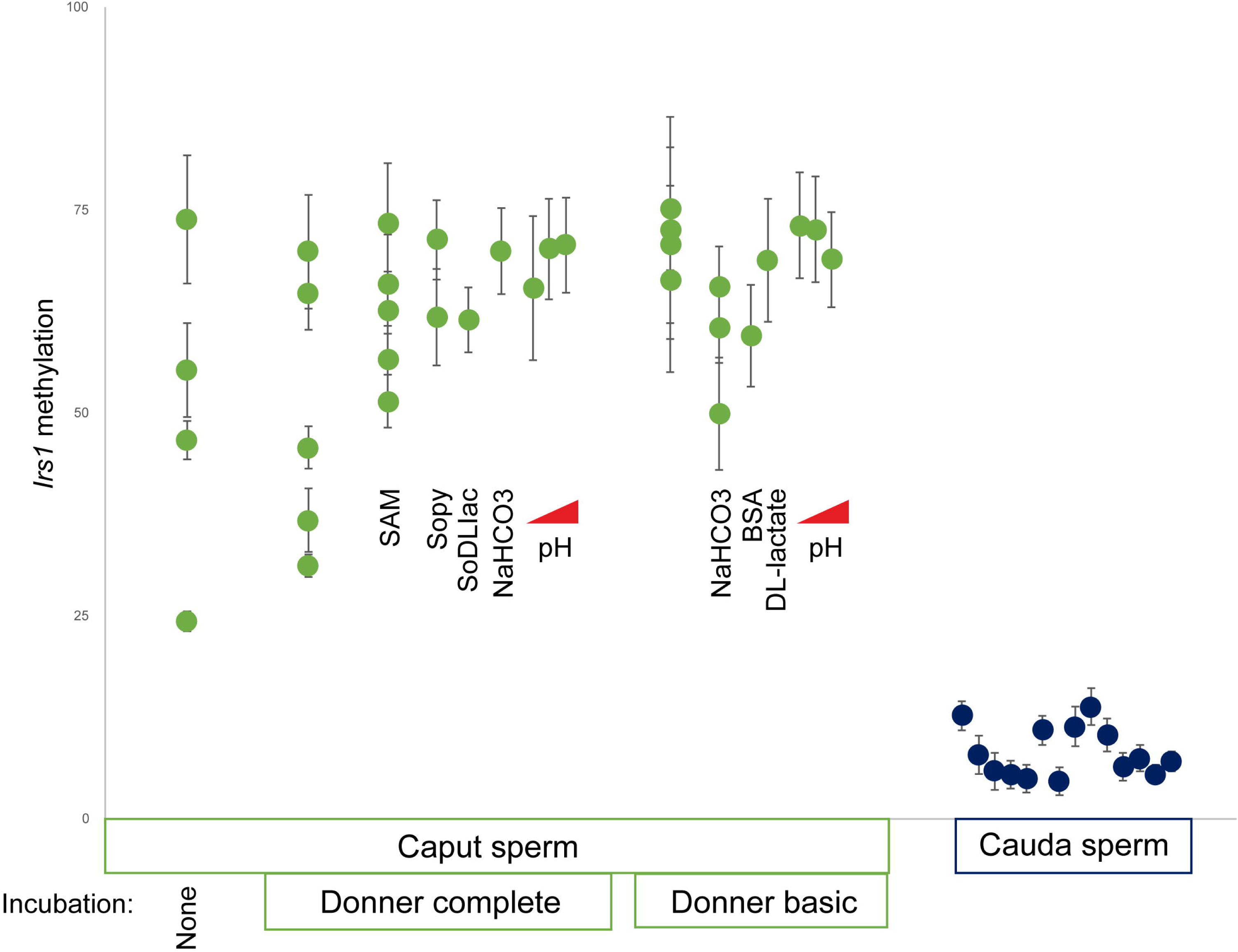
Caput sperm methylation status is stable under many different buffer conditions. Pyrosequencing data for caput and cauda sperm samples. Caput sperm samples were either processed for genomic DNA shortly after isolation, or were incubated for four hours at 37 C in various buffer conditions, as indicated. Buffer conditions are based on either Donners Basic (DB) or Donners Complete (DC), and were supplemented with various levels of NaHCO_3_, sodium pyruvate, sodium DL-Lactate, or adjusted to pH 6.5, 7.0, or 7.4 (red triangles). Although not indicated in the figure, cauda sperm samples (right) included samples subject to most of the buffer incubations used for caput sperm samples, none of which affected methylation in these samples.

**Figure 4-figure supplement 1.**
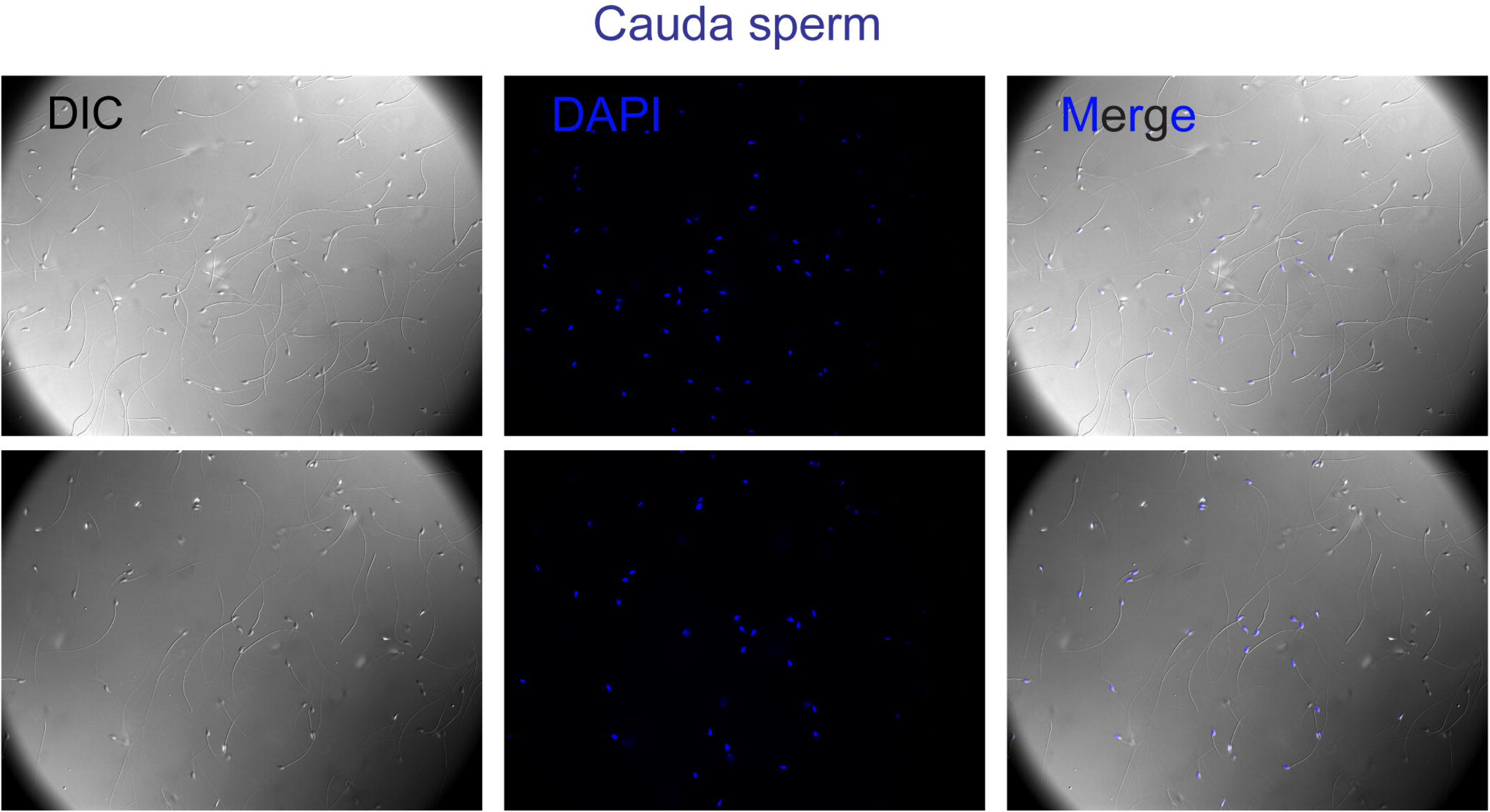
Absence of cell-free DNA from cauda sperm preps. Cauda epididymal sperm preparations were stained with DAPI as in **Figure 4A**. Here, DAPI staining was completely confined to sperm heads, and no webs of DNA were observed even at the highest exposures.

**Figure 5-figure supplement 1.**
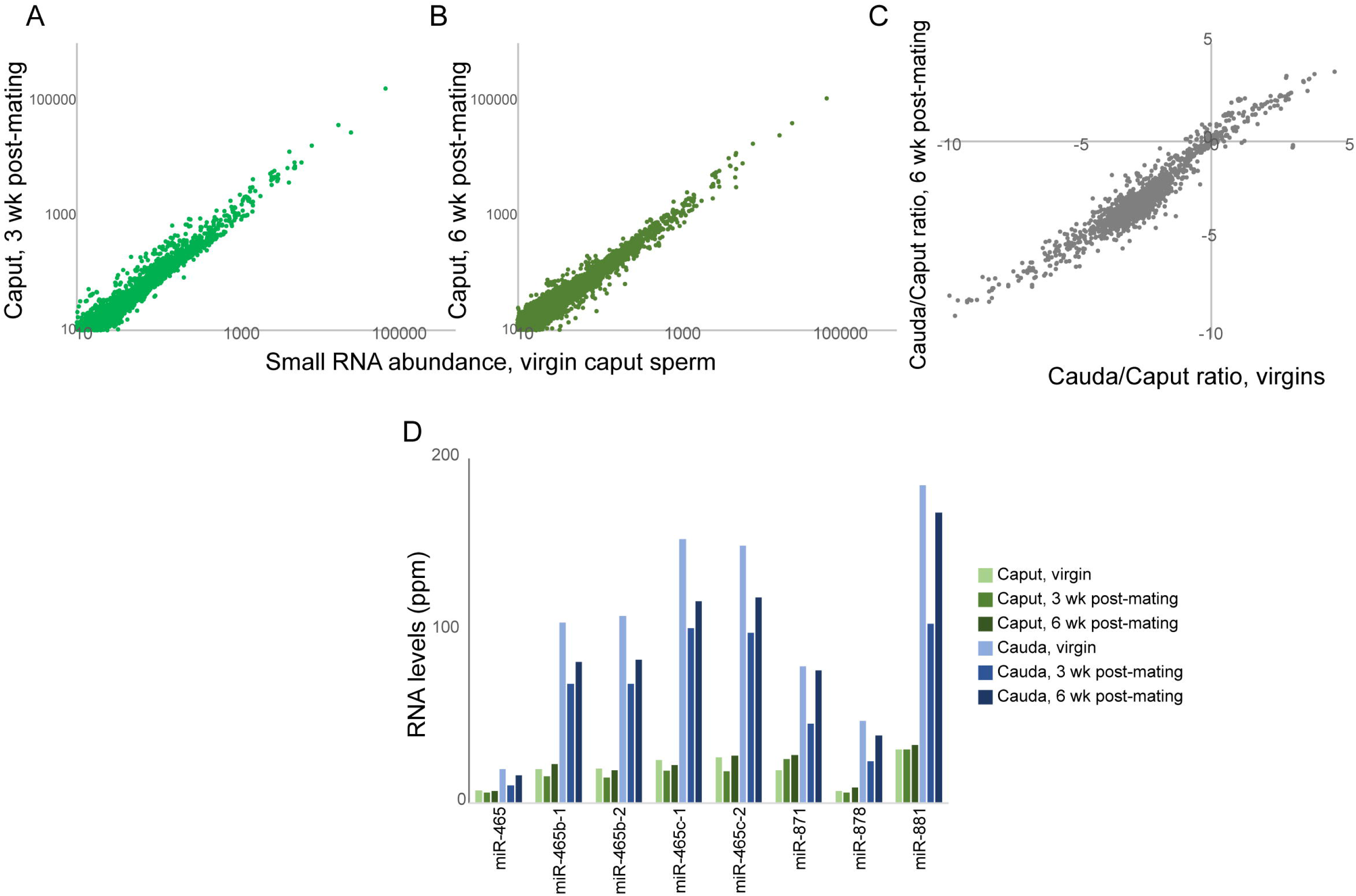
Mating does not impact the unusual small RNA payload of caput sperm. Several previous studies have documented significant differences between the RNA payload of caput and cauda sperm, including a dramatic loss of genomically-clustered microRNAs in caput sperm relative to both testicular and cauda sperm (Nixon et al., 2015; Sharma et al., 2016; Sharma et al., 2018). To determine whether this unusual small RNA profile was unique to caput sperm obtained from virgins, we obtained caput and cauda epididymal sperm from virgin males as well as males three and six weeks after successful mating, and characterized small RNAs by deep sequencing. A-B) Scatterplot compares small RNA levels in virgin caput sperm to levels in caput sperm 3 (A) or 6 (B) weeks after mating. Data are shown for all small RNAs with an abundance of at least 10 ppm in virgin caput sperm. C) Scatterplot shows enrichment/depletion in cauda sperm vs. caput sperm (calculated as log2(Cauda+1)/(Caput+1)) for virgin males (x axis) compared to males 6 weeks after mating (y axis). Overall enrichments for small RNAs in cauda or caput sperm remained highly correlated after mating. D) Example of cauda-enriched microRNAs from the X-linked miR-465 and miR-880 clusters. Enrichment in cauda sperm, and absence from caput sperm, was unaffected by mating status.

